# Set4 coordinates the activity of histone deacetylases and regulates stress-responsive gene expression within subtelomeric regions in yeast

**DOI:** 10.1101/2021.05.11.443697

**Authors:** Yogita Jethmalani, Khoa Tran, Deepika Jaiswal, Meagan Jezek, Mark Ramos, Shandon Amos, Eric Joshua Garcia, Maraki Y. Negesse, Winny Sun, DoHwan Park, Erin M. Green

## Abstract

The yeast chromatin protein Set4 is a member of the Set3-subfamily of SET domain proteins which play critical roles in the regulation of gene expression in diverse developmental and environmental contexts, although they appear to lack methyltransferase activity. The molecular functions of Set4 are relatively unexplored, likely due to its low abundance in standard growth conditions. We previously reported that Set4 promotes survival during oxidative stress and regulates expression of stress response genes via stress-dependent chromatin localization. In this study, global gene expression analysis and investigation of histone modification status has revealed a role for Set4 in maintaining gene repressive mechanisms within yeast subtelomeres under both normal and stress conditions. We show that Set4 works in a partially overlapping pathway to the SIR complex and the histone deacetylase Rpd3 to maintain proper levels of histone acetylation and expression of stress response genes encoded in subtelomeres. This role for Set4 is particularly critical for cells under hypoxic conditions, and the loss of Set4 decreases cell fitness and cell wall integrity in hypoxia. These findings uncover a new regulator of subtelomeric chromatin that is key to stress defense pathways and demonstrate a function for yeast Set4 in regulating repressive, heterochromatin-like environments.

## Introduction

The regulation of gene expression in response to changing environmental signals is dependent on a diverse set of chromatin-binding proteins, including transcription factors, histone-modifying enzymes, and chromatin remodeling complexes. Proteins containing a SET domain are well-established regulators of gene expression primarily through the catalysis of methylation of lysine residues within histones (1,2), though SET domain proteins also methylate non-histone substrates (3,4). A subfamily of SET domain proteins, often referred to as the Set3 subfamily, is characterized by divergent SET domains which appear to lack methyltransferase activity due to amino acid substitutions at key substrate binding interfaces (5,6). This subfamily includes the *Saccharomyces cerevisiae* paralogs Set3 and Set4, SET-9 and SET-26 from *Caenorhabditis elegans*, UpSET from *Drosophila melanogaster*, and the mammalian proteins MLL5 and SETD5 (7). Instead of directly catalyzing lysine methylation at chromatin, the Set3 subfamily of proteins are thought to regulate gene expression by binding to and regulating histone deacetylases (HDACs) at chromatin (8,9).

Our previous work identified a role for the yeast protein Set4, a paralog to Set3, in protecting cells during oxidative stress, primarily through gene expression regulation of stress response genes (10). Set4 is lowly expressed in yeast cells under standard laboratory growth conditions, although deletion of *SET4* increases sensitivity to acute oxidative stress and alters gene expression patterns (7,10-12), indicating a biological function for Set4 even at low abundance. Expression of *SET4* appears to be stress-regulated, as the transcript and protein levels increase in low oxygen, including hypoxic or anaerobic conditions (13,14). Other work has also implicated Set4 in the regulation of gene expression during hypoxia (14), where it was shown to repress ergosterol biosynthetic genes together with the transcriptional repressor Hap1 through the inhibition of the sterol-responsive activator Upc2 (14).

The subtelomeric regions of *S. cerevisiae* and other fungi, such as *Candida glabrata*, harbor many stress-response genes, particularly those that control adhesion, filamentation, and adaptation to anaerobic environments (15,16). Gene expression within subtelomeres is generally very low (17) due to regional silencing mechanisms, such as by the SIR histone deacetylase complex, and other transcriptional repressors (18). In other systems, Set3- and Set4-related proteins have been shown to maintain heterochromatic or repressive chromatin environments, including the fission yeast ortholog Set3 (19), the fly ortholog UpSET which interacts with the Rpd3/Sin3 deacetylase complex (20,21) and the *C. elegans* orthologs SET-9 and SET-26 which restrict spreading of H3K4me3-demarcated regions to regulate expression of germline-specific genes (22). While Set3 in budding yeast is critical for gene repression in multiple contexts, it is not known to have a specific role in maintaining silent chromatin states such as at subtelomeres, the mating type locus, or ribosomal DNA (rDNA) locus (8,9,23). Given the structural similarities between yeast Set3 and Set4 and orthologous proteins (7), we hypothesized that Set4 may function in regulating silent chromatin regions in yeast, especially since genes required for multiple stress response pathways are found within silent regions such as subtelomeres (18). Here, we demonstrate that Set4 calibrates gene expression within yeast subtelomeres under both normal and stress conditions and contributes to cell fitness and cell wall integrity in hypoxic conditions. In hypoxia, Set4 promotes subtelomeric chromatin binding of the HDACs Sir2 and Rpd3, which have previously been implicated in the regulation of stress response and subtelomeric genes (17,24-29). This maintains proper levels of histone acetylation and fine-tunes expression levels of genes within the subtelomere. These data uncover a key function of Set4 in controlling subtelomeric chromatin to coordinate proper gene expression in response to stress. Furthermore, our results indicate that while this regulatory role of Set4 is performed under non-stress conditions, it becomes critical for cells in response to certain environmental signals, including oxidative stress and limiting oxygen.

## Materials and Methods

### Yeast strains and growth conditions

The genotypes for all *Saccharomyces cerevisiae* strains used in this study are listed in **Table S1**. Strains carrying gene deletions were made using targeted PCR cassettes amplified from the pFA6a vector series (30). Double mutant strains were isolated following haploid mating, sporulation and tetrad dissection. All strain genotypes were confirmed by growth on the appropriate selective media and colony PCR using primers specific to individual gene deletions or epitope tag insertions. Standard media conditions for rich media (YPD; 1% yeast extract, 1% peptone, 2% dextrose) and synthetic complete (SC) or dropout media (US Biological) were used as necessary. For all growth assays, gene expression and chromatin immunoprecipitation experiments, yeast cultures were diluted and grown in appropriate media overnight to mid-log phase (OD_600_ ∼0.4-0.8) at 30°C. For hypoxic growth, the culture flasks were placed in BD GasPak EZ anaerobe pouch system and incubated at 30°C. For hydrogen peroxide-treated cultures, cells were grown to mid-log phase (OD_600_ ∼0.6-0.8) and then treated with 0.4mM H_2_O_2_ for 30 min (12).

### RNA Sequencing

Total RNA was extracted from yeast cells and subjected to Illumina-based RNA-sequencing at Genewiz (South Plainfield, NJ). Differential gene expression analysis was performed as previously described (31,32). Briefly, read quality control was analyzed using FastQC and adaptor removal and read trimming were performed with Trimommatic v.0.36 (33). Reads were mapped to the *S. cerevisiae* reference genome using the STAR aligner v.2.5.2b and Subread package v.1.5.2 was used for calculating gene hit counts (34). The data were normalized and log-fold change values were determined using DESeq2 (35). The raw and processed data for RNA-sequencing experiments are available on the Gene Expression Omnibus database at accession number GSE173901.

### Differential gene expression significance testing

For testing the significance of gene expression changes, we use a hybrid of two existing methods depending on the applicability of zero assumption in Efron (36): the center of the observed log_2_ fold-change (log FC) values consists of non-differentially expressed genes. One method is the local false discovery rate procedure (local FDR) which estimates the distribution of the non-differentially expressed genes based on zero assumptions instead of using the standard normal distribution. It can be a more powerful test when zero assumption holds: log FC values showed little change at most genes with small groups of up- and down-regulated genes exhibiting the most change (**Figure S1**). Local FDR analysis was employed to identify differentially-expressed genes at FDR ≤ 0.05 for datasets comparing wildtype and *set4*Δ cells. The false discovery rate is computed from these estimates and is controlled to be less than 5%. The method is implemented through the locfdr package in R (37).

The other method used for datasets comparing expression differences between aerobic and hypoxic conditions is the test procedure in DESeq2 after filtering absolute value of log FC > 1, the Wald test p-values which are adjusted (padjust) for multiple testing using the procedure of Benjamini and Hochberg (38). Since the test in DESeq2 does not need the structure assumption such as zero assumption, we applied it when there was large variability of log FC values and more of the non-differentially expressed genes spread out due to the discrepancy resulting from a larger number of up- and down-regulated genes. Padjust ≤ 0.05 are selected to be differentially expressed genes which represents FDR ≤ 0.05.

Gene ontology analysis was performed using the GO term function in Yeastmine and the GO term slim mapper through the Saccharomyces Genome Database. Telomere enrichment was determined by identifying the number of genes within 40 kb of the telomere end in each dataset analyzed and using a hypergeometric test to determine significance of enrichment and fold-enrichment over expected based on the total number of genes within subtelomeres in the genome.

### Gene expression analysis by quantitative reverse transcriptase PCR

Total RNA was extracted using 1.5 ml of mid log phase culture of yeast cells (OD_600_ ∼0.6-0.8) under different growth conditions. Masterpure Yeast RNA purification kit (Epicentre) was used to extract the RNA by following the manufacturer’s instructions. Turbo DNA-free kit (Ambion) was used to eliminate genomic DNA from the samples. cDNA was generated from 1 μg of total RNA using qMax cDNA synthesis kit (Accuris) containing both oligo dT and random hexamers for priming reverse transcription of mRNA. For quantitative PCR (qPCR) to check transcript levels 0.5 μl of cDNA was added to 1X qMax Green Low ROX qPCR mix (Accuris) with the appropriate gene specific primers (**Table S2**) in a 10 μl reaction. Real-time amplification was performed on a Bio-Rad CFX384 Real-time Detection System. Three technical replicates were performed for each reaction and a minimum of three biological replicates was performed for each experiment. Relative gene expression values were normalized to the control gene *TFC1*, whose expression has been shown to be stable under different growth conditions (39).

### Spot assays

For the telomere position effect spot assay, strains integrated with the *URA3* gene at *TELVIIL* were used (see **Table S1**; kindly provided by Paul Kaufman). Gene knockouts were created using insertion of targeted PCR cassettes amplified from the pFA6a vector series (30). Cells were grown overnight in YPD medium at 30°C and 0.1 OD units of the cultures were serially diluted and spotted on YPD plates (control) and 5-fluoroorotic acid (5-FOA) plates. The plates were observed and imaged for two days to analyze the growth pattern. For growth analysis of single and double mutant strains under aerobic and hypoxic conditions, yeast strains were grown overnight in YPD, diluted to OD_600_ ∼ 0.2 the next day, and grown to log phase. 0.1 OD_600_ units of the culture were serially diluted and spotted on YPD plates. For hypoxic conditions, the plates were incubated in BD GasPak EZ anaerobe pouches. The plates were observed and imaged for two days for aerobic conditions and eight days for hypoxic conditions.

### Telomere Southern Blot

Whole cell extract from wildtype and *set4Δ* strains was made by bead beating in phenol-chloroform-isoamyl alcohol. The extract was treated with RNase A and DNA was precipitated using ethanol. Genomic DNA was digested using the restriction enzyme XhoI, extracted with phenol-chloroform-isoamyl alcohol, and precipitated with ethanol. Digested DNA was subjected to electrophoresis on an 0.8% agarose gel in 0.5X TBE, the DNA was denatured in-gel and transferred onto a Hybond N+ nylon membrane (Amersham). The membrane was hybridized with a biotin-conjugated telomere probe (5’-biotin-CACACCCACACCCACACC-3’) and was imaged using a Chemiluminescent Nucleic Acid Detection Module Kit (Thermo Scientific) and a Li-Cor C-DiGit Chemiluminescent Western Blot scanner.

### Zymolyase sensitivity assay

WT (yEG001) and *set4*Δ (yEG322) were diluted and grown overnight to mid-log phase (OD_600_ ∼ 0.4-0.8) in aerobic or hypoxic conditions. Cells were collected and resuspended in 1 mL sorbitol buffer (1.2 M Sorbitol, 0.1 M KP04 pH 7.5) and 5 µl of 2-mercaptoethanol and 5 µl of 10 mg/mL 100T Zymolyase was added. Cells were incubated at room temperature, with occasional rocking, and the OD_600_ was measured every 5 minutes in 1% SDS. Time _50% OD_ was determined as the time elapsed for the cultures to reach 50% of the starting OD_600_, indicating 50% digestion by zymolyase and generation of spheroplasts.

### Trypan blue staining of cell walls

Detection of cell walls using Trypan blue was performed as described previously (40). Briefly, 1 ml of log phase (OD_600_ ∼ 0.6-0.8) culture under aerobic and hypoxic conditions was centrifuged at 10K rpm for 2 min. The cells were washed once in PBS and resuspended in 1 ml PBS. Trypan blue was added at a final concentration of 10 µg/ml. 5 µl of cells were observed on a slide with coverslip using a Leica SP5 confocal microscope. Image processing was performed using Image J. Staining intensity was determined by drawing a region around the stained area, measuring the mean gray value and subtracting the background signal for approximately 300 cells under aerobic conditions and 150 cells under hypoxic conditions.

### Chromatin Immunoprecipitation

Chromatin immunoprecipitation (chIP) was performed as described (41-43). Briefly, cultures were diluted and grown overnight to mid-log phase (OD_600_ ∼0.4-0.8). Cultures were then fixed with 1% formaldehyde for 20 minutes (histone chIPs) or 45 minutes (FLAG-Set4 and Sir3-HA chIPs). For the Rpd3-FLAG chIPs, a double crosslinking strategy was used as previously reported (44) to improve recovery of Rpd3-FLAG with chromatin. In this case, cells were harvested and resuspended in PBS. EGS (ethylene glycol bis (succinimidyl succinate)) was added to a final concentration of 1.5mM and cells were fixed for 30 min. Then 1% formaldehyde was added and cells were incubated for an additional 30 min. Quenching was performed with 0.5 M Tris-HCl pH 7.5 for 10 min. Cells were pelleted and washed 1X with TBS prior to lysis.

Whole cell extracts were made by bead beating and the chromatin was digested with micrococcal nuclease enzyme. The amount of chromatin used was 40 μg per IP (histone chIPs) and 100-300 μg per IP (FLAG-Set4, Sir3-HA, and Rpd3-FLAG chIPs). The antibodies were either pre-bound to protein A/G magnetic beads (Pierce) overnight (histone, Sir3-HA, Rpd3-FLAG chIPs) or pre-conjugated anti-FLAG M2 magnetic beads (Sigma-Aldrich) were used (FLAG-Set4 chIPs). The beads were added to the extracts and rotated overnight at 4°C. Protein-DNA complexes were eluted using 1% SDS and 0.1M NaHCO_3_, cross-links were reversed, and samples were treated with proteinase K and RNase A. DNA was extracted with phenol-chloroform-isoamyl alcohol and precipitated using ethanol. qPCR was performed as described above using 0.5 μl chIP DNA per reaction and gene-specific primers (**Table S2**). Three technical replicates were performed for each qPCR reaction and a minimum of three biological replicates were performed for each chIP experiment. Percent input was calculated relative to 10% of the input. The following antibodies were used for chIP: rabbit anti-H4K5ac (AbCam; catalog no. ab51997), rabbit anti-H4K12ac (EMD Millipore; catalog no. ABE532), rabbit anti-H4K16ac (EMD Millipore; catalog no. 07-329), rabbit anti-H3 (AbCam; catalog no. ab1791), rabbit anti-H3K4me3 (Active Motif; catalog no. 39159), rabbit anti-H3K9ac (EMD Millipore; catalog no. 06-942), mouse anti-FLAG (Sigma-Aldrich; catalog no. F1804), mouse anti-HA (EMD Millipore; catalog no. 05-904).

## Results

### Subtelomeric gene expression is disrupted in set4Δ mutants

To better define the contribution of Set4 to gene expression and any potential roles in silent chromatin regulation, we performed an RNA-sequencing experiment on wildtype and *set4Δ* cells in unstressed conditions (mid-log-phase growth, rich medium). Significantly differentially-expressed genes were identified based on log_2_ fold-change (log FC) in *set4Δ* cells relative to wildtype using local FDR ≤ 0.05 (see Materials and Methods; **Table S3**). In this analysis, 196 genes were identified as significantly differentially-expressed in *set4Δ* cells, with 75 genes up-regulated and 121 genes down-regulated in the absence of Set4 (**Figure 1A, Table 1**). We performed gene ontology analysis to identify enriched categories of genes among those differentially-expressed and identified no functional enrichment in genes down-regulated in *set4Δ* cells, though there is enrichment for genes involved in cell wall organization in those up-regulated in *set4Δ* cells (**Table 1**).

**Table 1.** Significant differentially-expressed genes in *set4Δ* mutants compared to wildtype under aerobic and hypoxic conditions.

|  | **Total genes** | **Up-regulated** | **Down-regulated** |
| --- | --- | --- | --- |
| **Aerobic** | 196 | 75<br>Cell wall organization ($9 \times 10^{-08}$) | 121<br>No enrichment |
| **Hypoxic** | 377 | 205<br>Cell wall organization ($3 \times 10^{-11}$) | 172<br>Cell wall organization ($2 \times 10^{-04}$)<br>DNA Integration ($2 \times 10^{-05}$) |

The number of significant differentially expressed genes in each category is indicated along with the enriched GO terms with *p* values indicated in parentheses.

**Figure 1:**
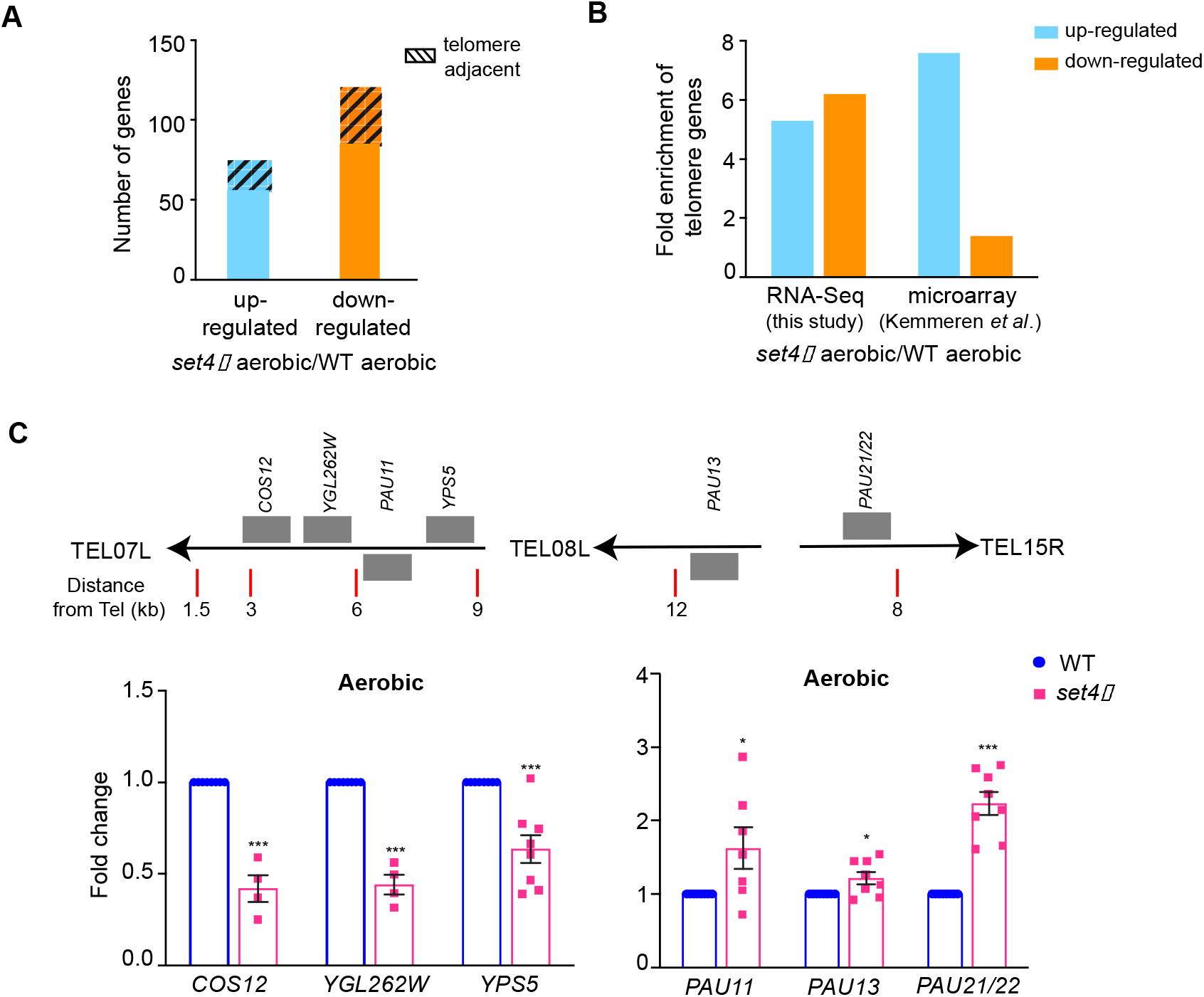
Set4 regulates the expression of subtelomeric genes. A. The total number of genes identified as up- or down-regulated (FDR < 0.05) from RNA-sequencing of *set4Δ* (yEG322) cells relative to wt (yEG001). Gene list provided in Table S3. The total number of telomere-enriched genes are indicated with the hashed box. B. The fold enrichment of differentially expressed subtelomeric genes (defined as less than 40kb from the chromosome end) in our RNA-sequencing data of *set4Δ* cells and in previously published microarray data (11). C. RT-qPCR of sub-telomeric genes from wt (yEG001) and *set4Δ* (yEG322) strains grown in YPD. Expression levels were normalized to *TFC1*. Fold change relative to wt is shown. The error bars represent S.E.M. from at least three biological replicates. Asterisks represent *p* values as calculated by an unpaired t test (* <0.05, **<0.01, *** <0.001).

We noted that many of the genes associated with cell wall organization are encoded within subtelomeric regions (29). Therefore, we next assessed the enrichment of genes within 40 kb of chromosome ends to determine whether there is a more general enrichment for differential expression of subtelomeric genes in *set4Δ* cells that is independent of gene functional category. We observed a more than five-fold enrichment in genes adjacent to telomeres (*p* = 1.60 × 10^−28^ for all genes; hypergeometric test; **Figure 1B**) within those differentially expressed in *set4Δ* mutants. We also analyzed previously published microarray data of *set4Δ* cells grown in synthetic medium (11). Interestingly, the differentially-expressed genes in this dataset also showed significant enrichment for subtelomeric genes (*p* value = 0.0003; **Figure 1B**), providing further evidence that Set4 may have a specific role in regulating expression of telomere-adjacent genes. In the same dataset (11), gene expression in *set3Δ* cells was also analyzed, which showed no significant enrichment for differential expression of subtelomeric genes (*p* value = 0.115). Together, these data suggest that under normal growth conditions, Set4 plays a specific role in regulating telomere-adjacent genes and genes linked to cell wall organization.

To further investigate these findings, we performed targeted gene expression analysis of wildtype and *set4Δ* cells using RT-qPCR. We investigated two classes of subtelomeric genes: (1) one set of genes (*COS12, YGL262W*, and *YPS5*) on the left arm of chromosome seven adjacent to *TEL07L*, a well-characterized site for chromatin-based silencing and telomere position effect; and (2), the seripauperin (*PAU*) genes, a highly homologous, subtelomeric gene family induced during different stresses–particularly anaerobic growth–that are thought to be important for cell wall remodeling or sterol uptake during stress (**Figure 1C**) (45,46). We monitored expression of *PAU11*, which is also located adjacent to *TEL07L*, and *PAU13*, using primers that uniquely amplify these genes (**Table S2**), as well as *PAU21* and *PAU22*, which have identical sequences (indicated as *PAU21/22* where appropriate). In the absence of Set4, *COS12, YGL262W and YPS5* were downregulated, whereas *PAU11, PAU13*, and *PAU21/22* were upregulated. These data suggest Set4 is important for both maintaining expression of some subtelomeric genes and repressing other subtelomeric genes under physiological, unstressed conditions.

The pattern of neighboring gene expression changes observed in *set4Δ* cells is consistent with a role for Set4 in altering regional chromatin structure. We therefore tested whether *set4Δ* cells showed any defects in a canonical telomere position effect (TPE) assay using a strain carrying *URA3* integrated near *TEL07L*. In this reporter assay, we did not observe any substantial change in *URA3* expression in the absence of Set4 or Set3, unlike the loss of silencing observed in *set1Δ* cells (**Figure S2A**). We also analyzed telomere length by Southern blot, which showed no difference between wildtype and *set4Δ* cells in the length of terminal telomere restriction fragments (**Figure S2B**). This indicates that Set4 has a specific regulatory role distinct from other TPE regulators and is not required for telomere length maintenance under normal conditions.

### *Deregulation of gene expression is enhanced in* set4Δ *mutants during stress*

In previous work, we demonstrated that Set4 promotes proper gene expression in response to oxidative stress (10). Many genes encoded within subtelomeres are stress response genes; therefore we analyzed whether some of these genes showed Set4-dependent changes in expression during oxidative stress. Upon treatment with hydrogen peroxide, genes that were downregulated in *set4Δ* cells did not show substantial change (*COS12* and *YGL262W*; **Figure S3A**). Genes that were up-regulated in *set4Δ* cells under normal conditions (e.g. *PAU13* and *PAU21/22*) were more highly upregulated in *set4Δ* cells in the presence of hydrogen peroxide, although repressed in wildtype cells, indicating that the loss of Set4 attenuates their repression in hydrogen peroxide.

Serratore et al. (14) previously showed that Set4 is important for the regulation of gene expression in hypoxic conditions and that hypoxia causes an increase in Set4 protein levels. As the *PAU* genes are highly upregulated during hypoxia (46), we tested their expression in *set4Δ* cells, along with the other telomere genes, under hypoxic conditions. We first tested our growth conditions for wildtype cells grown in aerobic and hypoxic conditions. We observed that we obtained the most consistent results by diluting stationary phase cultures to very low OD_600_ and allowing them to grow to OD_600_ ∼0.4-0.8 over the course of 18 hours in hypoxia, similar to how we tested *set4Δ* mutants sensitivity to oxidative stress (10,12). Under these conditions, the *PAU* genes were highly upregulated, and there was also significant upregulation of *YGL262W*, although expression of *COS12* and *YPS5* at *TEL07L* remained mostly unchanged in hypoxic compared to aerobic growth (**Figure 2A**). In the *set4Δ* strain grown under hypoxia, *COS12* and *YGL262W* expression showed no or minimal decrease in hypoxic conditions, whereas the expression of *PAU11, PAU13, PAU21/22*, and *YPS5* was significantly increased over the level of induction seen in wildtype cells (**Figure 2B**). These data indicate that the loss of Set4 leads to enhanced induction of hypoxia-regulated genes, including genes that are both negatively- and positively-regulated by Set4 under aerobic conditions (e.g., the *PAU* genes and *YPS5*, respectively). These observations parallel our findings in hydrogen peroxide treated cells, in which repression is inhibited at *PAU13, PAU21/22*, and *YPS5* (**Figure S3A**), indicative of a common gene regulatory role for Set4 under different stress conditions.

**Figure 2:**
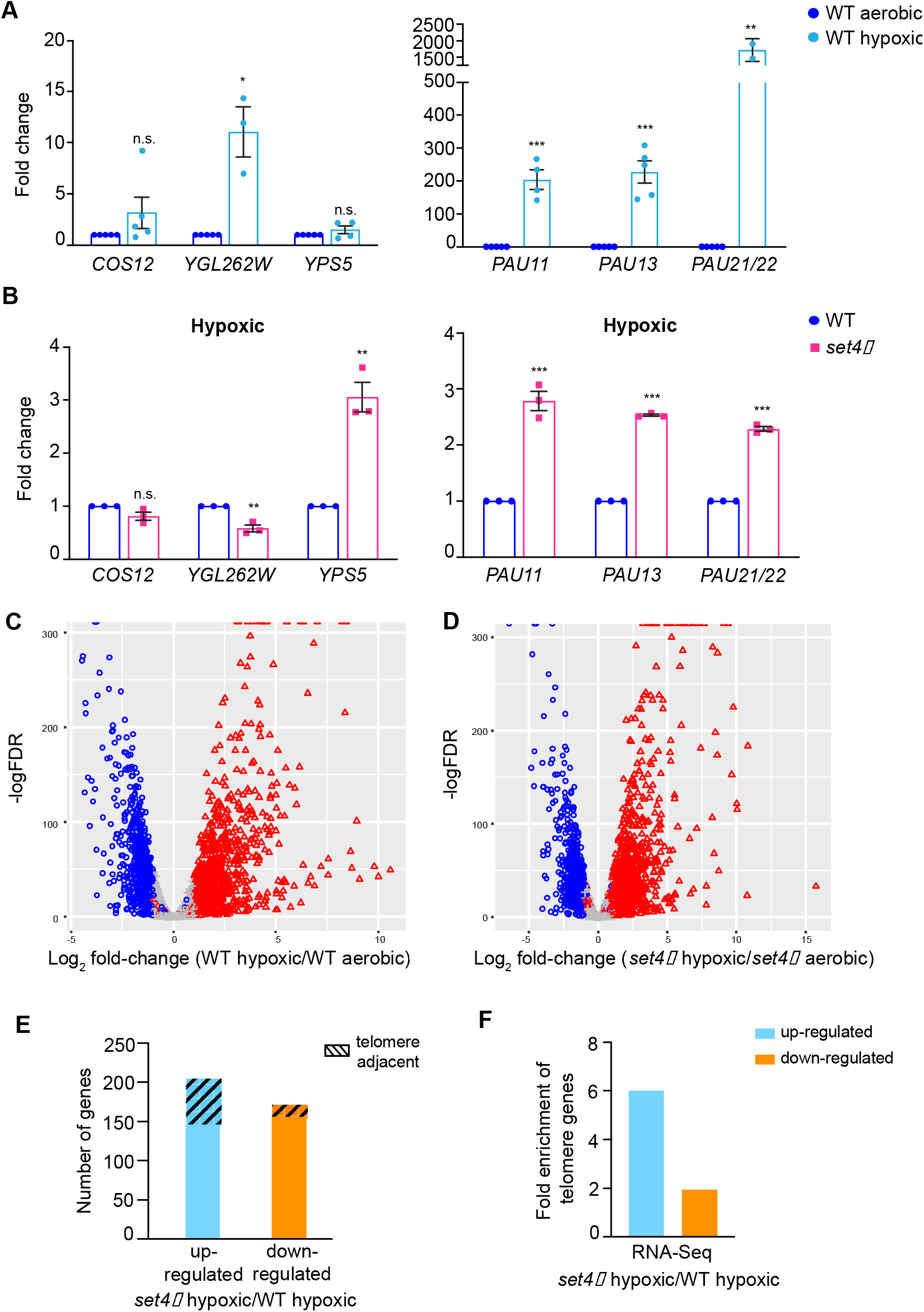
Stress response genes at subtelomeres are regulated by Set4. **A**. RT-qPCR of sub-telomeric genes from wt (yEG001) strains grown in YPD under aerobic or hypoxic conditions. Expression levels were normalized to *TFC1* and fold change relative to aerobic conditions is shown. **B**. RT-qPCR of subtelomeric genes from wt (yEG001) and *set4Δ* (yEG322) strains grown in hypoxia in YPD. Expression levels were normalized to *TFC1*. Fold change relative to wt in hypoxia is shown. For all panels, error bars represent S.E.M. from at least three biological replicates and asterisks represent *p* values as calculated by an unpaired t test (* <0.05, **<0.01, *** <0.001). **C, D**. Volcano plots depicting significantly differentially expressed genes (log FC ≥ 1.0, *p* ≤ 0.05) comparing wildtype hypoxic to wildtype aerobic cultures (**C**) and *set4Δ* hypoxic to *set4Δ* aerobic cultures (**D**). **E**. The total number of genes identified as up- or down-regulated (FDR < 0.05) from RNA-sequencing of *set4Δ* (yEG322) cells relative to wt (yEG001) in hypoxia. Gene list provided in **Table S3**. The total number of telomere-enriched genes are indicated with the hashed box. **F**. The fold enrichment of differentially expressed subtelomeric genes (defined as less than 40kb from the chromosome end) for those genes differentially-expressed between wildtype hypoxic cultures and *set4Δ* hypoxic cultures.

To address the role of Set4 in regulating subtelomeric genes more broadly during stress, we performed RNA-sequencing of wildtype and *set4Δ* cells grown under hypoxic conditions. In wildtype cells, growth in hypoxia induced widespread gene expression changes with 1056 genes up-regulated and 835 genes down-regulated (log FC ≥ 1.0, *p* ≤ 0.05; **Table S3, Figure 2C**). The significantly differentially-expressed genes encompassed a range of GO categories, including enrichment for genes associated with transmembrane transport, lipid metabolic process, and cell wall organization, among others, in the up-regulated genes (**Table S4**). The genes down-regulated in wildtype cells in hypoxia were highly-enriched for translation associated processes, mitotic cell cycle, cytoskeletal organization, cell wall organization, and lipid metabolic processes, among others (**Table S4**). The gene expression changes reported here are similar to those previously-described under hypoxic or anerobic growth of yeast (47,48).

In *set4Δ* cells, we observed a largely similar cohort of differentially expressed genes in hypoxia as in wildtype cells, with 1073 genes up-regulated and 917 genes down-regulated (log FC ≥ 1.0, *p* ≤ 0.05; **Table S3, Figure 2D**). These genes encompassed similar GO categories to those observed in wildtype cells (**Table S4**). There are more genes down-regulated in *set4Δ* cells grown in hypoxia compared to the total number of genes down-regulated in wildtype cells. These genes are distributed across a number of functional categories, including GO terms associated with translation-related processes, cytoskeletal organization, and DNA repair (**Table S4**).

When directly comparing wildtype and *set4Δ* cells in hypoxia, we identified 377 total genes differentially-expressed, with 205 genes up-regulated and 172 genes down-regulated in the absence of Set4 (**Table 1; Figure 2E**). Gene ontology analysis revealed enrichment for genes associated with cell wall organization in both the up- and down-regulated sets of genes and genes linked to DNA integration also enriched in the down-regulated genes (**Table 1**). Compared to aerobic conditions, this represents an increased number of cell wall organization genes misregulated in the absence of Set4. Interestingly, the down-regulated genes associated with the GO term DNA integration are almost entirely from Ty transposable elements (**Table S3**). Given that these are not differentially-expressed under aerobic conditions, this indicates enhanced repression of these genes without Set4 under hypoxia. In addition, previous work showed differential regulation of ergosterol biosynthetic genes in *set4Δ* mutants grown under hypoxia (14). However, we did not observe enrichment of ergosterol biosynthetic genes within the differentially-expressed gene set from our RNA-sequencing experiments (**Table S3**), nor by directly testing *ERG3* and *ERG11* expression using RT-qPCR (**Fig. S3B**). It is possible that differences in yeast strains or growth conditions, such as the time in hypoxia, may contribute to this difference in expression patterns.

Based on results obtained under aerobic conditions and RT-qPCR experiments performed on telomere genes, we predicted that genes with altered expression in *set4Δ* cells in hypoxia may show subtelomeric enrichment. Indeed, for those genes up-regulated in hypoxic *set4Δ* cells, there was six-fold enrichment for subtelomeric localization compared to expected (*p* = 2.79 × 10^−31^; hypergeometric test; **Figure 2F**) and almost two-fold enrichment for subtelomeric localization for down-regulated genes (*p* = 0.008). These data support our conclusions from the RT-qPCR experiments indicating enhanced expression changes in cell wall organization genes at subtelomeres in *set4Δ* cells under hypoxia and indicate a broad role for Set4 in regulating subtelomeric genes genome-wide under both normal and stress conditions.

### Set4 maintains cell wall integrity during hypoxic growth

Our previous work identified a role for Set4 in protecting cells during oxidative stress, likely through the regulation of gene expression. We showed that loss of Set4 increases sensitivity to oxidizing agents such as hydrogen peroxide, and Set4 overexpression increases survival upon hydrogen peroxide treatment (10). When growing cells under hypoxic conditions, we observed that *set4Δ* mutants grew more slowly and had smaller colony sizes (**Figure 3A**), indicating impaired growth under hypoxia. We hypothesized that the deregulated expression of the *PAU* genes, as well as other hypoxia-induced genes, may lead to disrupted cell wall integrity. We assayed cell growth in hypoxia with sorbitol to increase the osmolarity and suppress cell wall defects. As shown in **Figure 3A**, the growth differential between wildtype and *set4Δ* cells was decreased in the presence of sorbitol, suggesting stabilization of any cell wall defects in these cells. In addition, we tested the sensitivity of our strains to zymolyase digestion, which targets β-1,3 glucan linkages in the cell wall. As previously-shown (49), yeast grown under hypoxic conditions showed increased resistance to zymolyase compared to aerobic conditions (**Figure 3B**). However, *set4Δ* cells showed modestly more sensitivity to zymolyase digestion than wildtype cells in hypoxia, further indicating disrupted cell wall integrity.

**Figure 3:**
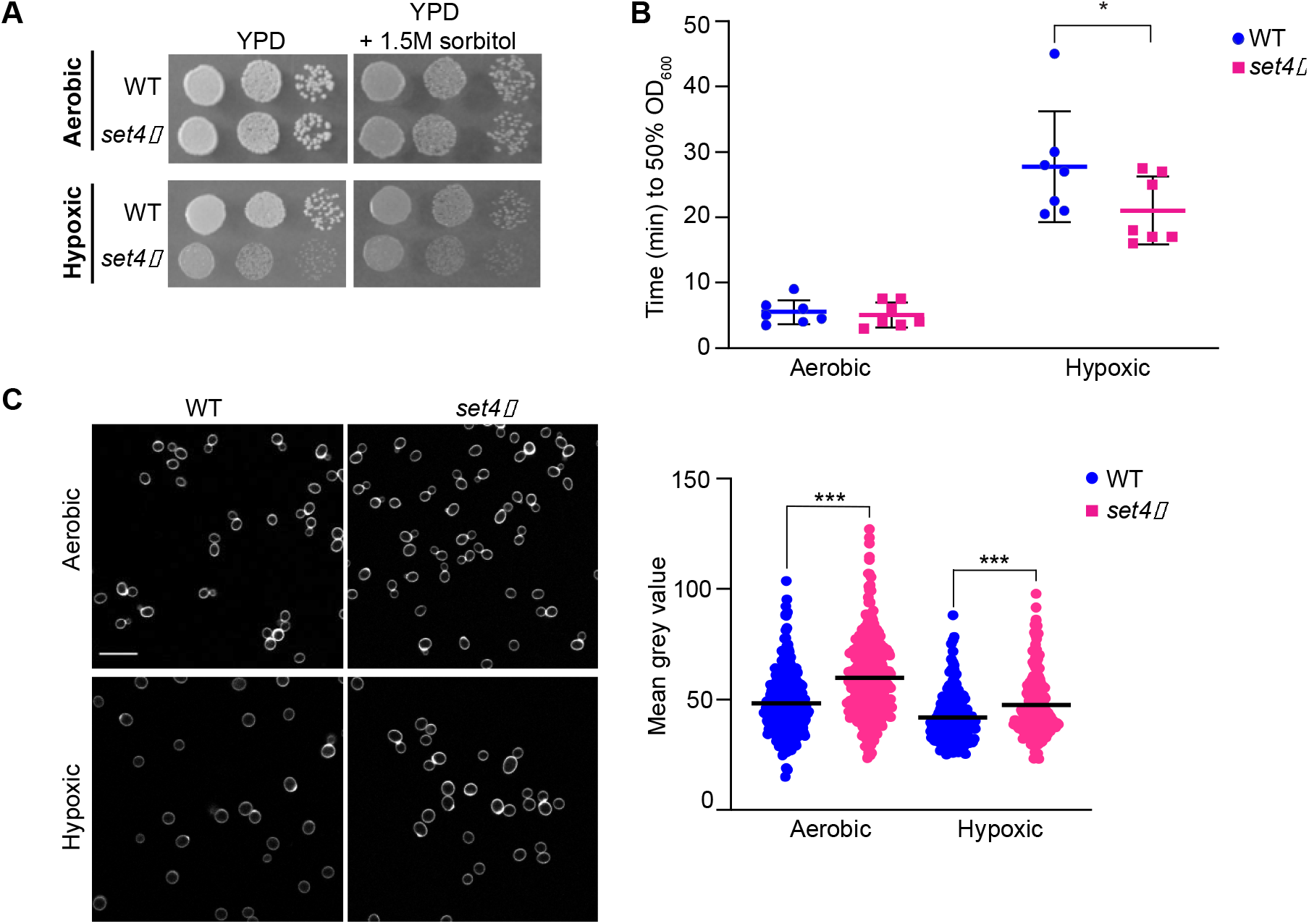
Set4 promotes cell fitness and cell wall integrity in hypoxia. **A**. Ten-fold serial dilutions of wt (yEG001) and *set4Δ* (yEG322) strains spotted on YPD or YPD with sorbitol and grown under aerobic (2 days) or hypoxic (8 days) conditions at 30°C. **B**. Scatter dot plot of the time elapsed for wt (yEG001) and *set4*Δ (yEG322) cultures grown in either aerobic or hypoxic conditions to reach 50% digestion by zymolyase. Error bars represent standard deviation (S.D.) from seven biological replicates. Asterisk represents *p* value as calculated by two-way ANOVA and Sidak’s multiple comparisons test (* <0.05). **C**. Fluorescence microscopy of wt (yEG001) and *set4Δ* (yEG322) cells grown under aerobic or hypoxic conditions and stained with Trypan blue. Quantitation of the mean gray value for approximately 300 cells under aerobic conditions and 150 cells under hypoxic conditions. Asterisk represents *p* value as calculated from an unpaired *t* test (*** <0.001).

Yeast cell walls show altered thickness and composition in hypoxia (49), which can be visualized using trypan blue, which stains yeast glucans and chitin (40). In *set4Δ* cells under aerobic conditions, we observed increased trypan blue staining of cell walls compared to wildtype (**Figure 3C**). In wildtype cells in hypoxia, chitin composition and cell wall mass decrease, lowering trypan blue staining (40). We observed an expected decrease in wildtype cells grown in hypoxia (**Figure 3C**), although staining remained relatively high in *set4Δ* cells in hypoxia, indicating an attenuation of this component of the hypoxic response. Altogether, these data are consistent with our previous findings of altered stress responses in the absence of Set4 (10).

### Set4 maintains histone acetylation levels at stress response genes within subtelomeric regions

Our data show deregulation of hypoxia response genes in the absence of Set4, particularly those located within subtelomeric regions and important for cell wall integrity. Multiple histone deacetylases (HDACs) have been shown to control expression of stress response genes within subtelomeric regions and are key regulators of the repressive chromatin environment at subtelomeres (26-29). Orthologs of Set4 in other organisms and the yeast protein Set3 are known to interact with or otherwise regulate the activity of HDACs in different chromatin environments (7,9,19,21,50,51). Thus, we hypothesized that Set4 may play a similar role at subtelomeric regions in yeast. To investigate this further, we tested the distribution of a series of acetylation marks previously implicated in the regulation of subtelomeric chromatin, including H4K5ac, H4K12ac, H4K16ac, H3K9ac, as well as the transcription-associated mark H3K4me3. We tested for association of Set4 at telomeric regions (*TEL07L*), as well as an internal site 12kb from the chromosome end which marks the approximate boundary with euchromatin (*TEL07L*_*boundary*_). We also tested localization at the promoters of the *PAU* genes in wildtype and *set4Δ* cells. In aerobic conditions, we observed lower levels of histone acetylation close to the telomere (*TEL07L* primer set), and increased acetylation levels and H3K4me3 at more distal regions such as the *TEL07L* boundary region (**Figure 4A**). This is the expected distribution pattern of histone acetylation and H3K4me3 at subtelomeres and provides both positive and negative controls for the chIP.

**Figure 4:**
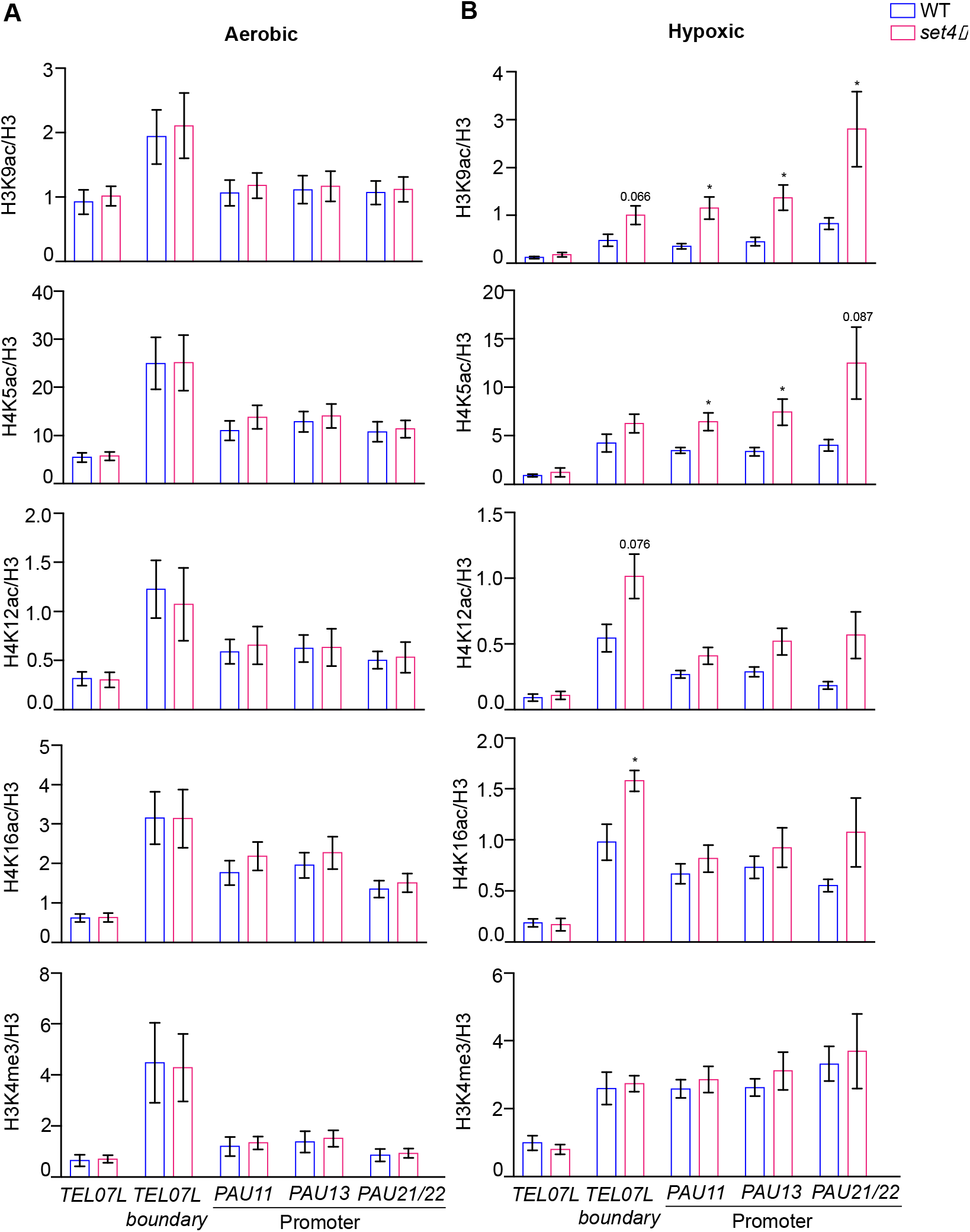
Histone acetylation increases at subtelomeric chromatin in *set4Δ* cells in hypoxia. chIP of H3K9ac, H4K5ac, H4K12ac, H4K16ac, and H3K4me3 at subtelomeric regions from wt (yEG001) and *set4Δ* (yEG322) strains grown to mid-log phase in YPD under aerobic (**A**) or hypoxic (**B**) conditions. Percent input of each acetyl or methyl mark are shown relative to percent input of total H3 levels. A minimum of three biological replicates are shown for histone mark chIPs and six biological replicates of histone H3 chIP was performed. The histone H3 immunoprecipitation is more efficient and consistent than histone H4, and therefore was used to normalize all of the histone modifications tested to total histone levels. For all panels, error bars indicate S.E.M. and asterisks represent *p* values as calculated by unpaired t tests (* <0.05, **<0.01, *** <0.001).

In the absence of Set4, there was relatively little change in the abundance of these marks at any of the regions tested under aerobic growth. However, in hypoxic conditions, we observed increased acetylation, particularly at telomere-distal locations and the promoters of the *PAU* genes (**Figure 4B**). The largest increase in acetylation was observed for H3K9ac, although H4K16ac and H4K5ac also showed marked increases at subtelomeric regions in *set4Δ* cells. The overall abundance of histone acetyl marks was not changed in *set4Δ* cells, although we did observe a global decrease in H4K16ac in hypoxic conditions compared to aerobic conditions (**Figure S4A**). We also note that there was no change in H3K4me3 levels in *set4Δ* cells in aerobic or hypoxic conditions (**Figure 4A-4B**), indicating that this mark is not regulated by Set4 at subtelomeric regions. Combined, these findings demonstrate increased acetylation at multiple histone sites upon loss of Set4 in hypoxic conditions, consistent with our observations of enhanced activation of the *PAU* genes and less repression of other subtelomeric genes (e.g. *COS12, YGL262W*) in *set4Δ* cells.

### Disrupted localization of HDACs at subtelomeric regions in set4Δ mutants

The HDACs Sir2 and Rpd3 are both known regulators of silent chromatin near telomeres (17,18,27,52) and have also been implicated in the regulation of stress response genes, including those induced during hypoxic or anaerobic growth (26,28,29,53). These observations, and our findings of altered levels of histone acetylation in the absence of Set4, led us to test the hypothesis that Set4 works with HDACs to maintain telomeric chromatin structure. We investigated the distribution of Rpd3 and the SIR complex in wildtype and *set4Δ* cells under hypoxic conditions. We focused on hypoxia as both the gene expression data and histone modification chIP data suggest a much larger dependence on Set4 in hypoxic than aerobic conditions.

Chromatin immunoprecipitation of epitope-tagged Sir3, which is the direct chromatin-binding component of the SIR protein complex, was performed in *set4Δ* cells under hypoxic conditions. We observed the expected occupancy of Sir3-HA primarily near telomeric chromatin at *TEL07L*, as well as secondary localization at the promoters of *PAU* genes, as previously demonstrated in aerobic conditions (28). In hypoxia, Sir3-HA localization at telomeric chromatin decreased in *set4Δ* cells relative to wildtype (**Figure 5A**), suggesting that Set4 promotes the proper association of the SIR complex with telomeres in these conditions. In agreement with previous findings (28), we observed more binding of Sir3 to *PAU13* and *PAU11* promoters compared to *PAU21/22* promoters, indicating that *PAU13* and *PAU11* may be more dependent on the SIR complex for maintaining repression. These findings are also consistent with the increase in acetylation in the region observed in hypoxia (**Figure 4**), including H4K16ac, the primary substrate of Sir2. While we did not observe any differences in protein expression levels of Sir3-HA between wildtype and *set4Δ* cells, nor between aerobic and hypoxic conditions (**Figure S4B**), the increased chromatin association of Sir3 in hypoxia in wildtype cells (**Figure S5A**), which is consistent with the global decrease in H4K16ac levels observed in hypoxia-treated cells (**Figure S4A**).

**Figure 5:**
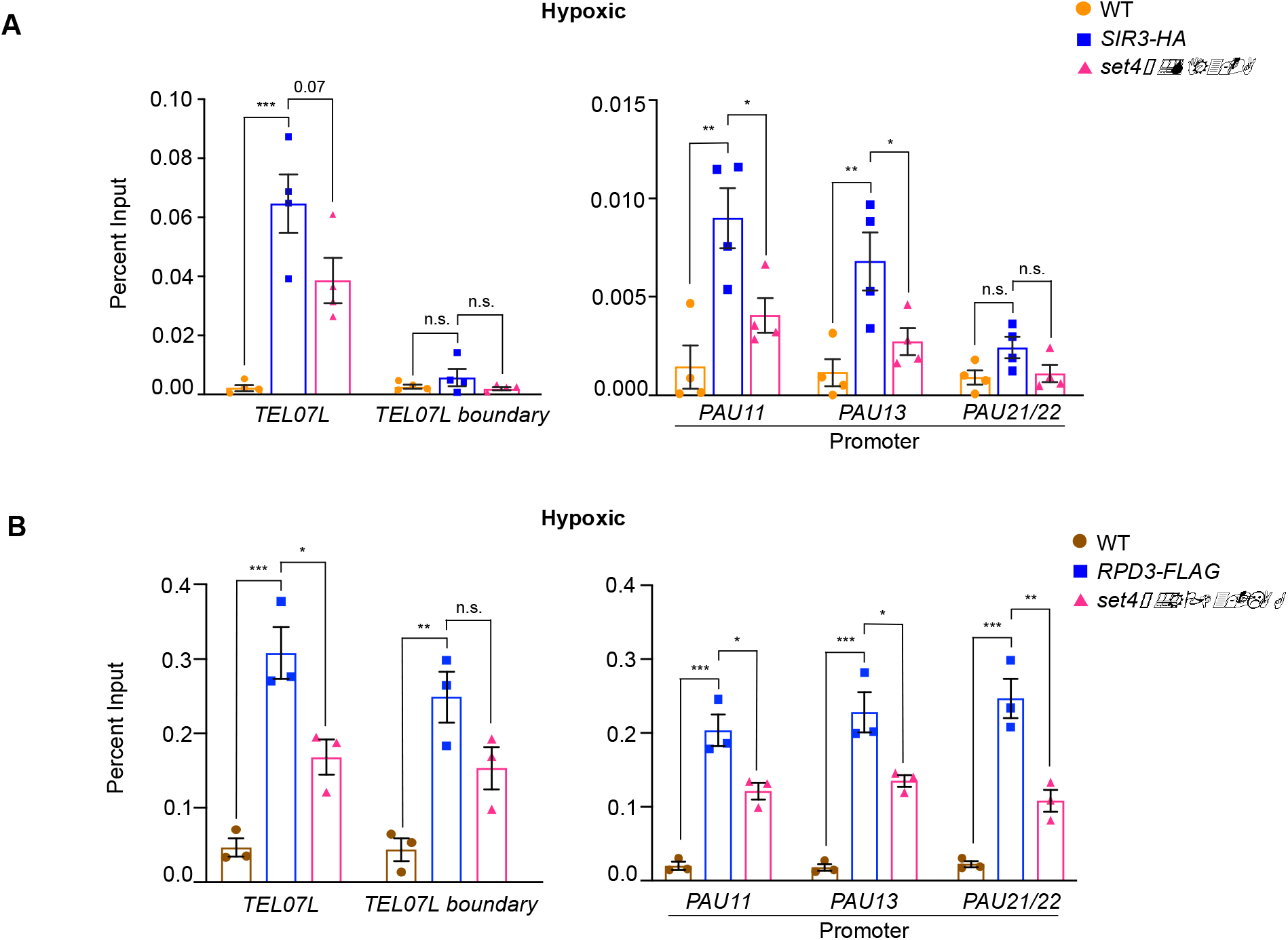
Disrupted HDAC distribution at subtelomeric chromatin in the absence of Set4. **A**. chIP of wt (yEG001), Sir3-HA (yEG873) and *set4Δ* Sir3-HA (yEG874) strains grown to mid-log phase in YPD in hypoxic conditions. **B**. chIP of wt (yEG001), Rpd3-FLAG (yEG956) and Rpd3-FLAG *set4Δ* (yEG1010) strains grown to mid-log phase in YPD in hypoxic conditions. For both panels, percent input from at least three biological replicates is shown. The error bars indicate S.E.M. and asterisks represent *p* values as calculated by one-way ANOVA and Tukey’s post-hoc test (* <0.05, **<0.01, *** <0.001).

We similarly evaluated the distribution of Rpd3-FLAG at subtelomeres in the absence of Set4. In hypoxic conditions, Rpd3-FLAG showed decreased binding in the absence of Set4 (**Figure 5B)** indicating that Set4 promotes the localization of Rpd3-FLAG to subtelomeric regions. There were no changes in total Rpd3-FLAG protein levels in *set4Δ* mutants (**Figure S4C**). These observations are consistent with increased acetylation within the region in *set4Δ* cells (**Figure 4**) and the altered gene expression patterns observed in *set4Δ* cells.

To further define the interaction of Set4 with the Sir2 and Rpd3 HDACs in regulating gene expression, we generated double mutant strains carrying *set4Δ* and *rpd3Δ* or *sir2Δ* and monitored gene expression changes and cell growth in hypoxia. Both *set4Δ* and *rpd3Δ* cells showed growth defects in hypoxic conditions compared to aerobic conditions, although there was no clear defect in *sir2Δ* cells and *sir2Δ set4Δ* cells grew very similarly to *set4Δ* single mutants (**Figure S5B**). The loss of Rpd3 resulted in a severe growth defect in hypoxia (**Figure S5C**), and *rpd3Δ set4Δ* cells grew similarly to *rpd3Δ* single mutants, suggesting that Rpd3 and Set4 may contribute to a shared pathway regulating growth in hypoxia.

We next evaluated gene expression at subtelomeres in these single and double mutant strains. As expected, we observed de-repression of telomere-adjacent genes in *sir2Δ* cells under aerobic conditions (**Figure S5D**). In contrast, we observed enhanced repression of telomere-adjacent genes *COS12, YGL262W*, and *YPS5*, as well as lower expression of two out of three *PAU* genes tested, *PAU11* and *PAU13*, in *rpd3Δ* cells (**Figure S5D**). These data are consistent with previous reports indicating an anti-silencing function for Rpd3 at subtelomeres (27).

In *sir2Δ* cells grown in hypoxic conditions, *COS12* and *YGL262W* were de-repressed, as expected (**Figure 6A**). Similar expression was observed in the *sir2Δ set4Δ* mutants, indicating that expression levels of these genes are largely regulated by the SIR complex in hypoxia. However, we observed a different expression pattern of genes that show enhanced activation in *set4Δ* cells in hypoxia, including *YPS5* and the *PAU* genes. These genes showed increased repression in *sir2Δ* cells compared to wildtype in hypoxia, however this repression was alleviated in the *sir2Δ set4Δ* double mutants. *PAU11* and *PAU13* were expressed at a similar level in the double mutant as wildtype cells, whereas *PAU21/22* and *YPS5* showed slightly higher expression than in wildtype. These data indicate an antagonistic function of the SIR complex and Set4 in balancing the expression of hypoxia-induced genes.

**Figure 6:**
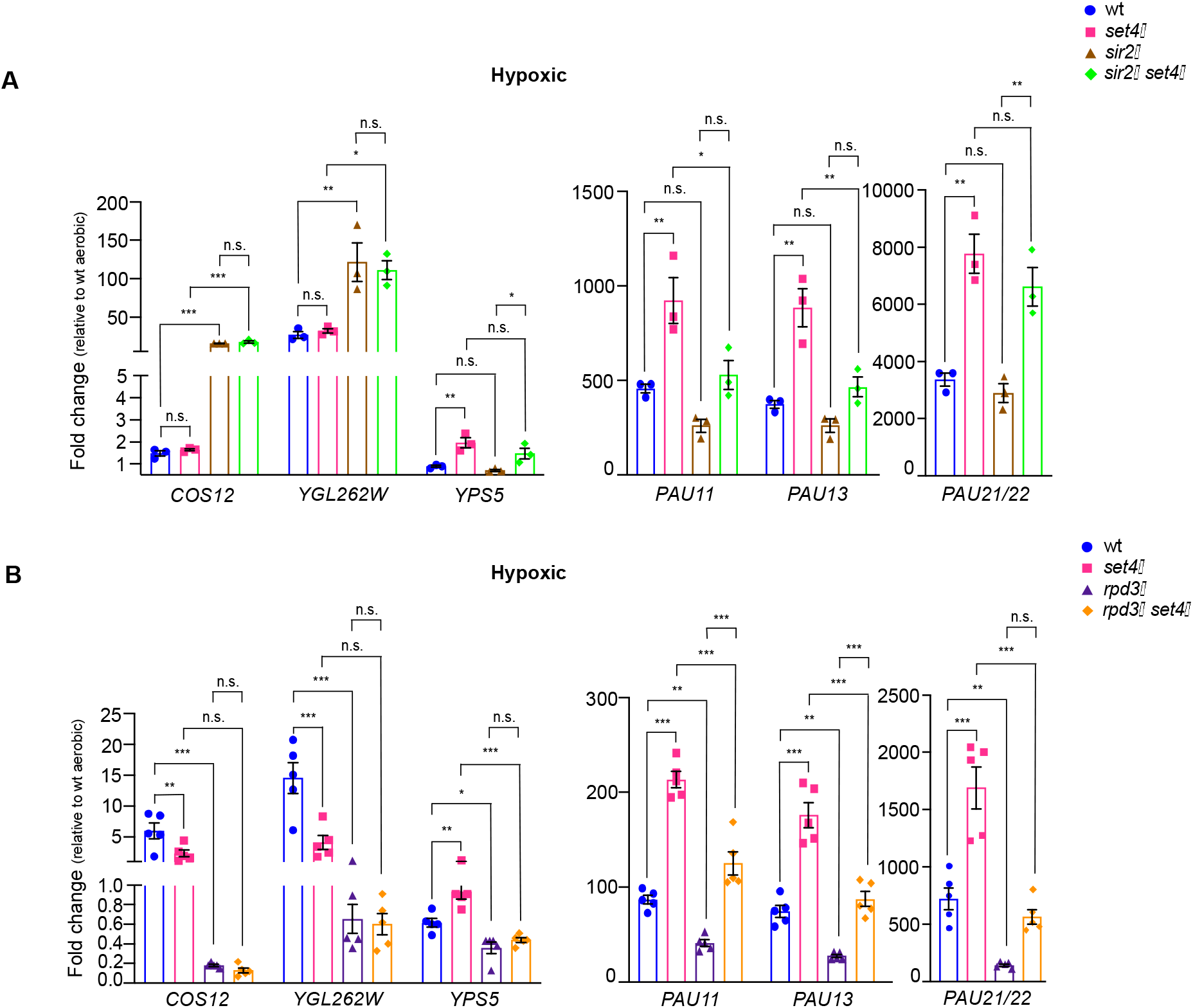
Genetic interactions of Set4 with HDACs Sir2 and Rpd3 in regulating subtelomeric gene expression during stress. **A**. RT-qPCR of subtelomeric genes from wt (yEG001), *set4Δ* (yEG1004), *sir2Δ* (yEG1002) and *set4Δ sir2Δ* (yEG1005) strains grown under hypoxic conditions in YPD. **B**. RT-qPCR of subtelomeric genes from wt (yEG919), *set4Δ* (yEG920), *rpd3Δ* (yEG921) and *set4Δ rpd3Δ* (yEG922) strains grown under hypoxic conditions in YPD. For all experiments, expression levels were normalized to *TFC1*. Error bars represent S.E.M. from at least three biological replicates. For all panels, asterisks represent *p* values as calculated by one-way ANOVA and Tukey’s post-hoc test (* <0.05, **<0.01, *** <0.001).

We also investigated changes in gene expression in the *rpd3Δ set4Δ* strain under hypoxic conditions. Rpd3 has been directly implicated in regulating the expression of genes induced during anaerobic growth (26). In the absence of Rpd3, repression of all of the subtelomeric genes remained largely intact compared to the induction observed in wildtype and *set4Δ* cells in hypoxic conditions. However, in the *rpd3Δ set4Δ* cells, repression of the *PAU* genes was relieved and induction closer to wildtype expression levels was observed (**Figure 6B**). The telomere-adjacent genes *COS12* and *YGL262W* showed similar expression levels in *rpd3Δ* and *rpd3Δ set4Δ* cells, indicating that loss of Set4 was not sufficient to overcome repression of these genes in the absence of Rpd3. These data suggest that, similar to its interaction with the SIR complex, Set4 counterbalances Rpd3 function in regulating expression of the *PAU* genes (and likely other genes induced in limiting oxygen). However, this is a gene-specific interaction, as Set4 and Rpd3 appear to function independently at other telomere adjacent genes such as *COS12, YGL262W* and *YPS5*.

### Set4 localizes to subtelomeric chromatin in hypoxia

Previous work from our lab showed that Set4 is a chromatin-associated protein and localizes to the promoters of genes that are induced during oxidative stress, particularly in the presence of stress (10). Another report has also shown that Set4 localizes to promoters of ergosterol biosynthetic genes during hypoxia (14). To investigate the localization of Set4 at subtelomeres and whether the changes in subtelomeric gene expression and acetylation levels are due to local occupancy by Set4, we performed chromatin immunoprecipitation (chIP) under both aerobic and hypoxic conditions using a strain expressing N-terminally FLAG-tagged Set4 from its endogenous locus and monitored binding to *TEL07L* and *TEL07L*_*boundary*_ regions and at the promoters of *PAU* genes. In aerobic conditions, we did not detect significant association of Set4 at any of these regions (**Figure S6**). However, under hypoxia, we observed binding of Set4 to a region near *TEL07L* as well as the promoters of the *PAU* genes (**Figure 7**). This binding was enhanced relative to a negative control region at *CENVX*. In addition, we also tested the promoters of other non-telomeric genes known to be regulated by Set4 during stress (*CTT1, PNC1, ERG3*, and *ERG11*) (10,14). Set4 localized to the promoters of *CTT1* and *PNC1*, as expected based on our previous findings (10), and was highly enriched at *ERG3* and *ERG11* gene promoter (**Figure 7**). These results are consistent with a previous report showing binding to *ERG3* and *ERG11* promoters in hypoxia (14), however our data indicate that these are not a major site for gene regulation by Set4 under similar conditions (**Figure S3B**).

**Figure 7:**
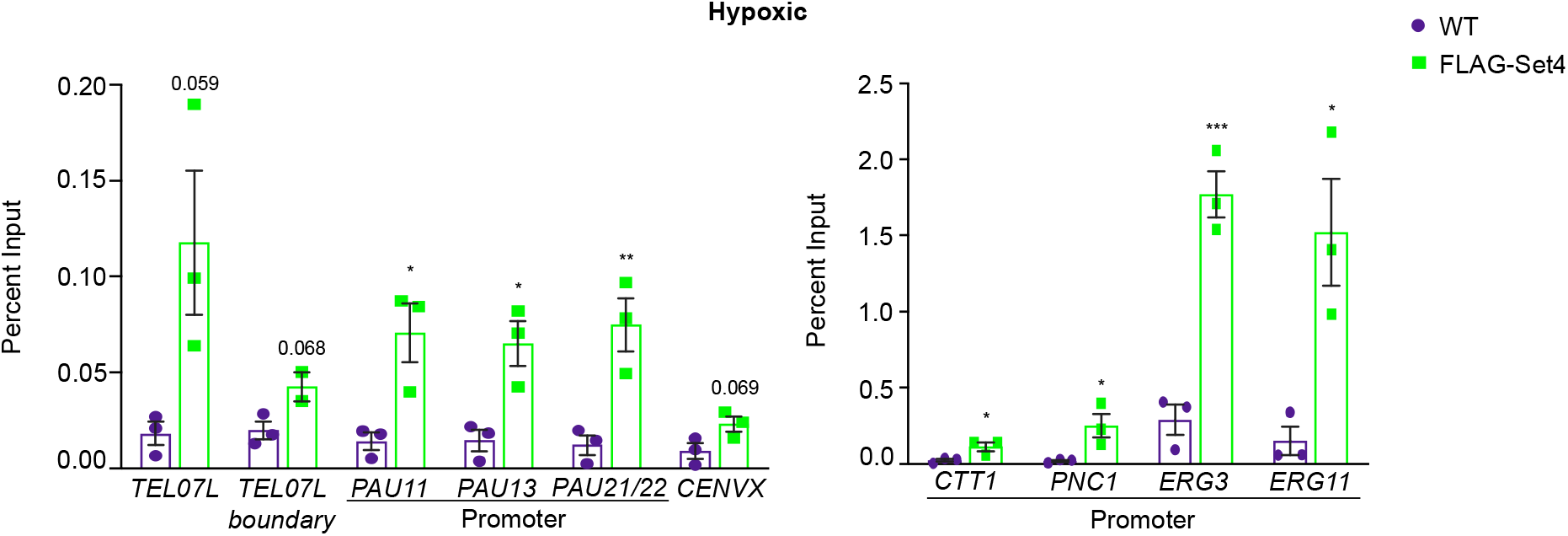
Set4 localizes to subtelomeric chromatin during stress. chIP of FLAG-Set4 from cells grown under hypoxic conditions. Percent input from three biological replicates is shown. Left graph shows regions tested here for histone acetylation levels and HDAC binding. Right graph shows promoter regions previously identified as binding locations in under stress conditions (10,14). Error bars represent S.E.M. and asterisks represent *p* values as calculated by an unpaired t test (* <0.05, **<0.01, *** <0.001).

The binding of Set4 to subtelomeric chromatin under stress suggests that Set4 may directly influence gene expression within the region. Set4 expression increases dramatically during hypoxia (14), therefore we expect increased association with chromatin under these conditions. In aerobic conditions, the abundance of Set4 is very low (10), which likely limits our ability to detect it by chIP. Combined with our gene expression data, we expect that Set4 may be present at subtelomeric chromatin at levels below the limit of detection in aerobic conditions, and Set4 abundance and localization near telomeres increases in hypoxia.

## Discussion

Yeast subtelomeres are enriched for stress response genes, and proteins orthologous to Set4 are known regulators of heterochromatin and gene silencing (19-22). Previous studies have highlighted a role for Set4 as a calibrator of stress-responsive gene expression (10,14). Here, we uncovered a role for Set4 in regulating genes within the repressed subtelomeric regions of budding yeast under both normal and stress conditions, particularly during hypoxia. Gene expression and chromatin immunoprecipitation analysis indicate that Set4 works together with the SIR complex and Rpd3 within the subtelomeres to fine-tune expression levels of stress-response genes. The loss of Set4 also decreases survival and cell wall integrity in hypoxia. Therefore, Set4 helps to maintain the proper balance of expression of stress response genes to promote survival during stress.

### Set4-dependent regulation of subtelomeric gene expression under both normal and stress conditions

Previous work has shown that Set4 localizes to the promoters of oxidative stress-induced genes following hydrogen peroxide treatment (10) and ergosterol biosynthetic genes during hypoxia (14). Set4 is lowly expressed under normal conditions, and its localization to these promoters was only detected during stress. We also observed enrichment of Set4 within subtelomeric regions, specifically during hypoxia, when Set4 protein abundance is dramatically increased (14). Changes in gene expression of telomere-adjacent genes and the stress-induced *PAU* genes were observed under both normal and stress conditions; however, the dependence on Set4 was clearly enhanced during stress. RNA-seq revealed an overall increase in differential gene expression between wildtype and *set4Δ* cells in hypoxia, although we note that there was little change in the pattern of gene expression changes observed compared to aerobic conditions (GO analysis and genomic location).

Consistent with changes in gene expression, there were greater changes in histone acetylation levels in hypoxia compared to normal conditions in *set4Δ* cells. We postulate that Set4 is present within subtelomeres (and likely other chromatin regions) even under normal, unstressed conditions, as we observe Set4-dependent changes in gene expression; however, the standard chIP assay is not sufficiently sensitive to detect this low abundance protein. In hypoxic conditions, the differences in gene expression and histone acetylation in *set4Δ* cells compared to wildtype cells are exacerbated, and we observe a clear localization of Set4 to subtelomeric chromatin. The increased abundance of Set4 in hypoxia (14) allows us to readily detect the protein using chIP. Combined with our previous results showing increased chromatin association of Set4 during oxidative stress (10), these data indicate that the gene regulatory role for Set4 is more critical during stress. This suggests that one component of the cellular response to certain types of stress is to increase Set4 protein levels and/or increase its association with chromatin to promote stress-responsive gene expression programs. Currently, this role for Set4 has only been linked to oxidative stress and limiting oxygen (hypoxic or anaerobic) conditions. It remains to be determined whether or not Set4 is a general stress response factor, similar to the Msn2 and Msn4 transcription factors (54), or if it has a specialized role under certain types of stress.

### Set4 coordinates histone deacetylases to regulate subtelomeric chromatin structure

The chromatin structure at subtelomeric regions of *S. cerevisiae* is maintained by multiple HDACs to generate a hypoacetylated state, which keeps gene expression levels low (17,18). Members of the Set3 subfamily of SET-domain proteins, including Set3, UpSET and SETD5 are all known to physically interact with histone deacetylases (7), and loss of function of these proteins leads to aberrantly high levels of histone acetylation (8,21,51). Protein-protein interaction analysis under hypoxic conditions revealed interactions of Set4 with other chromatin regulators, though not HDAC complex members (14), however further analysis may reveal how Set4 influences HDAC function. Using chromatin immunoprecipitation, we observed decreased binding of both the SIR complex and Rpd3 within subtelomeric regions in cells lacking Set4 under hypoxic conditions, when Set4 expression is high. Not all Rpd3 or SIR protein binding is lost in the absence of Set4, suggesting other targeting mechanisms of both HDACs are still intact. However, the reduced presence of each of these HDACs is consistent with increases in local histone acetylation in *set4Δ* cells in hypoxia (**Figure 8**). In addition, gene expression analysis in *sir2Δ set4Δ* and *rpd3Δ set4Δ* double mutants indicated that the repression of the *PAU* genes observed in the absence of either HDAC alone is relieved upon loss of Set4. At subtelomeric chromatin, Rpd3 has been reported to antagonize the spread of the SIR complex in silent chromatin regions (17,27,52). Our data suggests a model in which the loss of Set4 diminishes the ability of either Sir2 or Rpd3 to fine-tune *PAU* gene expression levels, causing aberrantly high activation of these genes. Despite the antagonism between the SIR complex and Rpd3, the reduction in both of their levels in *set4Δ* mutants likely reduces aberrant spreading of the HDACs, allowing *PAU* gene expression to approach or exceed wildtype levels in *set4Δ sir2Δ* and *set4Δ rpd3Δ* mutants.

**Figure 8:**
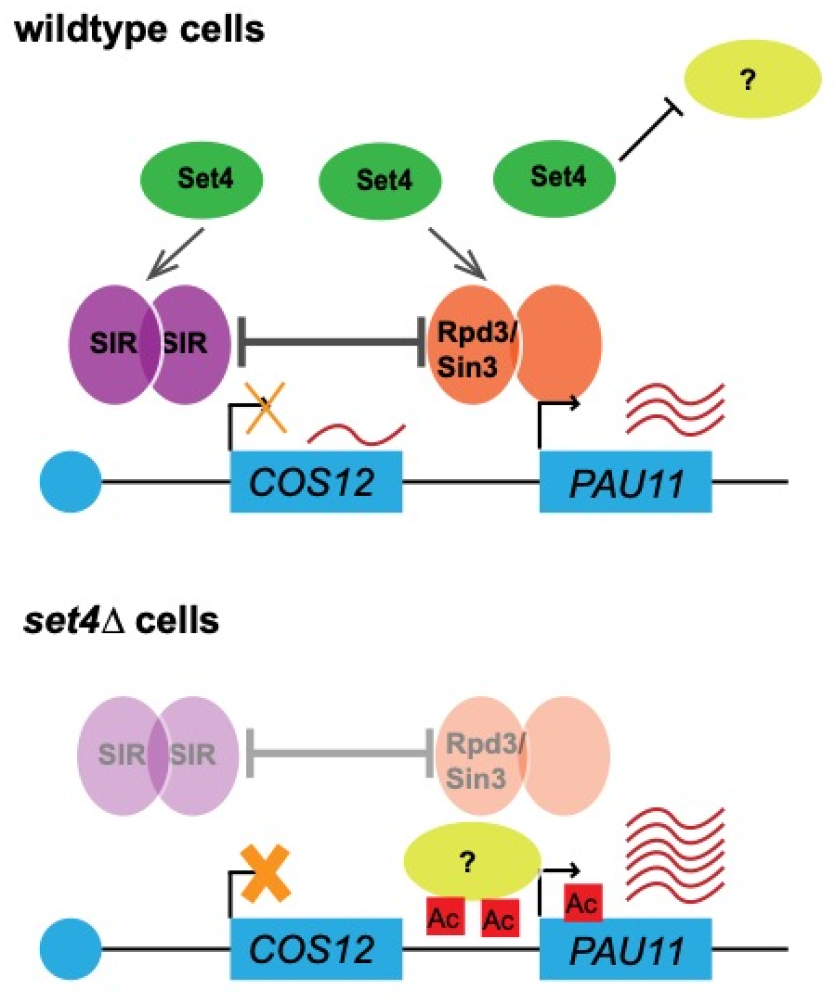
Model for Set4 function in subtelomeric gene regulation during hypoxic stress. A partial depiction of *TEL07L* is shown indicating genes subject to telomere position effect (TPE) silencing, such as *COS12*, and genes repressed under standard growth conditions and induced in stress, such as *PAU11*. In wildtype cells, Set4 promotes the association of the SIR complex (Sir2/3/4) and Rpd3 with subtelomeric chromatin. The presence of these HDACs represses telomere-adjacent genes subject to TPE, such as *COS12*, and genes induced in limiting oxygen, such as *PAU11*. Set4, either alone or in cooperation with the SIR complex, Rpd3, or other yet unidentified chromatin regulators, may also inhibit the binding or activity of factors important for the positive regulation of stress response genes at subtelomeres (indicated by a question mark). In the absence of Set4, both the SIR complex and Rpd3 binding are diminished, resulting in increased histone acetylation and enhanced activation of *PAU11* (and other *PAU*) genes. Genes subject to TPE, such as *COS12*, show enhanced repression upon loss of Set4, possibly due to diminishment of the antagonism between Rpd3 and the SIR complex, or due to compensation by other HDACs when Rpd3 and Sir2 levels are disrupted (56). This role for Set4 is most critical during stress, such as hypoxia when Set4 levels increase and it localizes to subtelomeric chromatin. Further genetic and physical interaction studies of Set4 at chromatin are likely to define the additional factors functioning with Set4, Rpd3 and the SIR complex in fine-tuning stress response genes within yeast subtelomeres.

It is also feasible that, in addition to maintaining proper SIR complex and Rpd3 levels at subtelomeres, Set4 works alone or in cooperation with these HDACs to inhibit association of a positive regulator of hypoxia-induced genes. Previous work has demonstrated that Rpd3 promotes the association of the transcription factor Upc2 with some anaerobic response genes (26). Upc2 is required for the induction of the *PAU* genes and works with the SAGA transcriptional activator and histone acetyltransferase complex to promote *PAU* gene expression in hypoxia (55). Interestingly, it has been reported that Set4 blocks Upc2 activity at ergosterol biosynthetic genes, thereby repressing these genes in hypoxia (14). Determining the interaction of Set4 with Upc2 and Rpd3 at the *PAU* genes may shed further light on regulatory mechanisms controlling their expression in hypoxia.

Altogether, this study provides new insights into the types of genes regulated by Set4 and the chromatin-based mechanisms through which it acts, as well as identifies a new telomere regulator in stress conditions. We have identified a role for Set4 in maintaining heterochromatic structures in yeast, which aligns its functions with metazoan orthologs previously implicated in heterochromatin maintenance (20-22), and expands our understanding of the role for Set4 during stress. Our data indicating decreased fitness and cell wall integrity of cells lacking Set4 in hypoxic conditions support the conclusion that Set4 promotes cell survival during stress, which is consistent with our previous findings identifying a role for Set4 in protecting cells during oxidative stress (10). Additional studies of Set4, and other Set3-related proteins, are likely to further our understanding of gene regulatory mechanisms and chromatin-mediated stress defense pathways.

## Data availability

The raw and processed data for RNA-sequencing experiments is available on the Gene Expression Omnibus database at accession number GSE173901.

## Acknowledgements

The authors thank all members of the Green lab for helpful discussions, technical assistance, and feedback on the manuscript. We thank Dr. Paul Kaufman for yeast strains and Dr. Philip Farabaugh for sharing equipment. We are also grateful to Tagide deCarvalho of the Keith R. Porter Imaging Facility for assistance with microscopy.

