## Supplemental Figures and Tables for "Set4 coordinates the activity of histone deacetylases and regulates stress-responsive gene expression within subtelomeric regions in yeast"

**Supplemental Material**

Table S1. Yeast strains used in this study

| Strain | Background | Genotype | Source |
| --- | --- | --- | --- |
| yEG001 | BY4741 | *MATa his3Δ1 leu2Δ0 met15Δ0 ura3Δ0* | YKO |
| yEG322 | BY4741 | *MATa set4Δ::HIS3MX* | (1) |
| yEG513 | BY4741 | *MATa set4::FLAG-SET4* | (1) |
| yEG873 | BY4741 | *MATa SIR3-HA::NATMX* | This study |
| yEG874 | BY4741 | *MATa set4Δ::HIS3MX SIR3-HA::NATMX* | This study |
| yEG956 | BY4741 | *MATa RPD3-FLAG::KANMX* | This study |
| yEG1010 | BY4741 | *MATa set4Δ::HIS3MX RPD3-FLAG::KANMX* | This study |
| yEG917 | BY4742 | *MATα sir2Δ::KANMX* | This study |
| yEG997 | BY4741 | *MATa set4Δ::HIS3MX sir2Δ::NAT3MX* | This study |
| yEG919 | BY4741 | *MATa his3Δ1 leu2Δ0 met15Δ0 ura3Δ0* | This study |
| yEG920 | BY4741 | *MATa set4Δ::HIS3MX* | This study |
| yEG921 | BY4742 | *MATα rpd3Δ::KANMX* | This study |
| yEG922 | BY4742 | *MATα rpd3Δ::KANMX set4Δ::HISMX* | This study |
| yEG017 | W303 | *MATa leu2-3,112 ura3-1 his3-11,15 trp1-1 ade2-1 can1-100 URA3-VIIL* | Paul Kaufman |
| yEG910 | W303 | *MATa set4Δ::KANMX URA3-VIIL* | This study |
| yEG909 | W303 | *MATa set3Δ::KANMX URA3-VIIL* | This study |
| yEG392 | W303 | *MATa leu2-3,112 ura3-1 his3-11,15 trp1-1 ade2-1 can1-100 URA3-VIIL set1Δ::HIS3MX* | This study |
| yEG952 | BY4742 | *MATα his3Δ1 leu2Δ0 met15Δ0 ura3Δ0* | This study |
| yEG953 | BY4742 | *MATα set4Δ::HIS3MX* | This study |
| yEG954 | BY4741 | *MATa hst1Δ::KANMX* | This study |
| yEG955 | BY4741 | *MATa hst1Δ::KANMX set4Δ::HISMX* | This study |

Table S2. Oligos used in this study

| Gene | Analysis | Sequence |
| --- | --- | --- |
| *COS12* | RT-qPCR | 5’-CATTACAAATACTCCGGGTATAGACA-3’  5’-GCAGCTGGAACCATCAAAA-3’ |
| *YGL262W* | RT-qPCR | 5’-GAGAATTACTCTGACATTGGAGATGA-3’  5’-TTGTCATTACAGAAGCCATCAAC-3’ |
| *YPS5* | RT-qPCR | 5’-CCTCCACAAACGGTGTACCT-3’  5’-TTGCAATAGGCAATGTCAGC-3’ |
| *PAU11* | RT-qPCR | 5’-CGCAACCACCACTCTAGCTC-3’  5’-TGAGCCAAGTGAGCTCTGAT-3’ |
| *PAU13* | RT-qPCR | 5’-TGAAACCTACCCGGTTGAAG-3’  5’-GGGGCAATACCAGTCAACAT-3’ |
| *PAU21/22** | RT-qPCR | 5’-CTTGGTCGAATTGGGTGTTT-3’  5’-TCTGTTGGATGAGCTGCTTG-3’ |
| *ERG11* | RT-qPCR | 5’-CCTCTTATTCCGTCGGTGAA-3’  5’-TGTGTCTACCACCACCGAAA-3’ |
| *TEL07L* | chIP | 5’-AGCCCGAGCCTGTACTAAAT-3’  5’-CAAAAGAAACTTTTCATGGCA-3’ |
| *TEL07L boundary* | chIP | 5’-AGCCATGCGGAAGTTATTTT-3’  5’-TCGACAATAAATAACGCATCG-3’ |
| *PAU11 promoter* | chIP | 5’-CGGGTATAAATAGAGCTGCTTCA-3’  5’-TGCTGGTATAAGCTTAACAGGAAAG-3’ |
| *PAU13 promoter* | chIP | 5’-GATGACTGATGAAGGCATGG-3’  5’-GCTTAACAGGAAGGGAAGGAA-3’ |
| *PAU21/22* promoter* | chIP | 5’-GTGATCATGAAGTTGTGGGAAA-3’  5’-CGATTCGTTAACAGATGCTCCT-3’ |
| *CTT1 promoter* | chIP | 5’-ATTCGACGTAGCCTGGACAC-3’  5’-TGGAATAGAGGTAAAGCAACGA-3’ |
| *PNC1 promoter* | chIP | 5’-TTCAAGGGGCAGGGGTTT-3’  5’-TATTAGCACATCATAATCGTATCTGGA-3’ |
| *ERG3 promoter* | chIP | 5’-CCGATGGCTGCGATAAACGA-3’  5’-TCGCTGCTGAACCTCTTGTT-3’ |
| *ERG11 promoter* | chIP | 5’-TTGCCGGGTTGGACAATCTT-3’  5’-TCGTTTCGTTTAGGGCCAGC-3’ |
| *CENVX* | chIP | 5’- TGCTTTCATAATACCCCACGA-3’  5’- AGGCAAAGGACGCACATATCT-3’ |

**PAU21/22* primers recognize both *PAU21* and *PAU22* due to repetitive sequence.

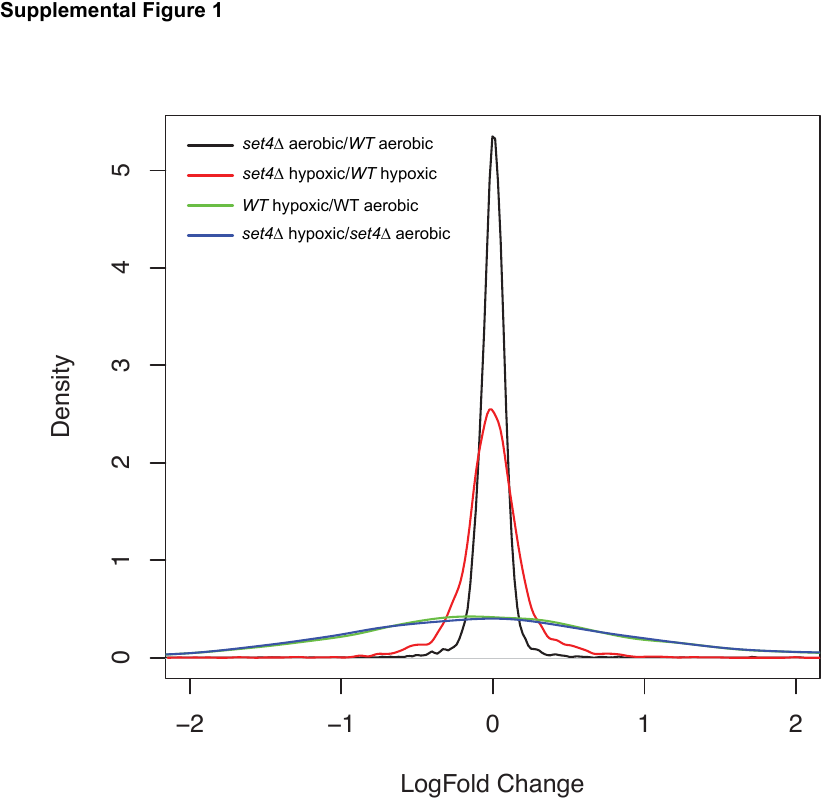

**Figure S1: Distribution of log_2_ fold-change for RNA-sequencing datasets.**

Smoothened histogram representing log_2_ fold-change values for RNA-sequencing experiments comparing *set4Δ* aerobic/WT aerobic, *set4Δ* hypoxic/WT hypoxic, WT hypoxic/WT aerobic, and *set4Δ* hypoxic/*set4Δ* aerobic.

**Figure S2: Set4 is not implicated in canonical telomere position effect (TPE) or telomere length regulation.**

**A.** Ten-fold serial dilutions of wt (yEG001), *set1Δ* (yEG392), *set4Δ* (yEG910) and *set3Δ* (yEG909) carrying a *URA3-VIIL* reporter were spotted on YPD and SC-URA + 5-FOA plates to assay telomere position effect. **B.** Southern blot using a telomere probe of two isolates of wt (yEG001) and *set4Δ* (yEG322) strains cultured in YPD.

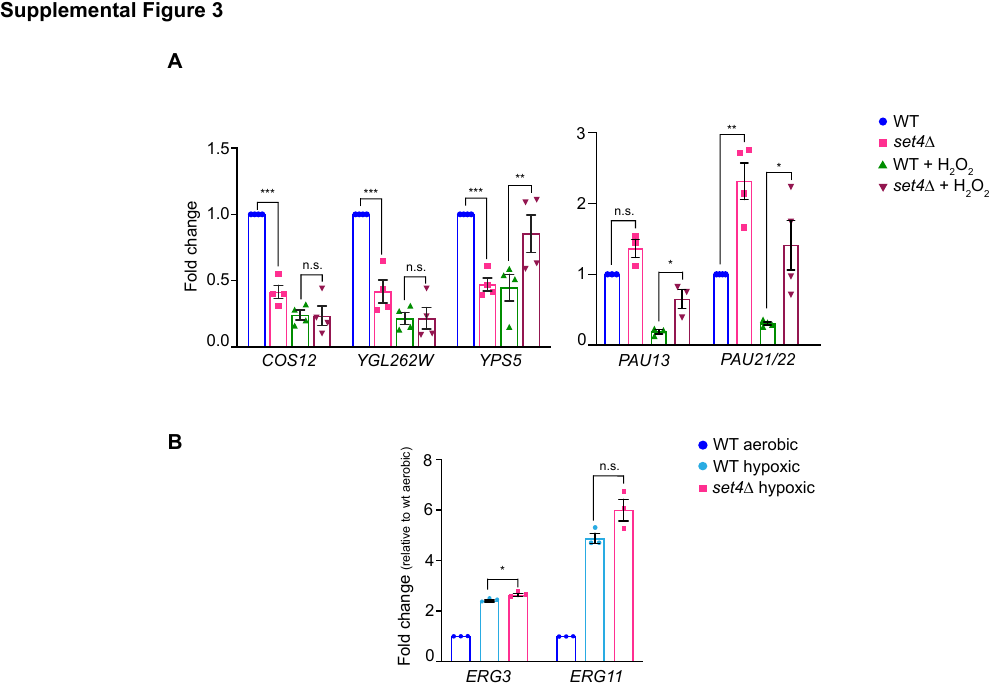

**Figure S3: Subtelomeric gene regulation in oxidative stress and ergosterol biosynthetic gene regulation in hypoxia.**

**A**. RT-qPCR of sub-telomeric genes from wt (yEG001) and *set4Δ* (yEG322) strains grown in YPD and treated with 0.4mM H_2_O_2_ for 30 min. Expression levels were normalized to *SCR1*. Fold change relative to wt without H_2_O_2_ treatment is shown. **B**. RT-qPCR of *ERG3* and *ERG11* from wt (yEG001) and *set4Δ* (yEG322) strains grown in YPD under aerobic or hypoxic conditions. Expression levels were normalized to *TFC1*. Fold change relative to wildtype aerobic is shown. For all panels, error bars represent S.E.M. from at least three biological replicates and asterisks represent *p* values as calculated by an unpaired t test (* <0.05, **<0.01, *** <0.001).

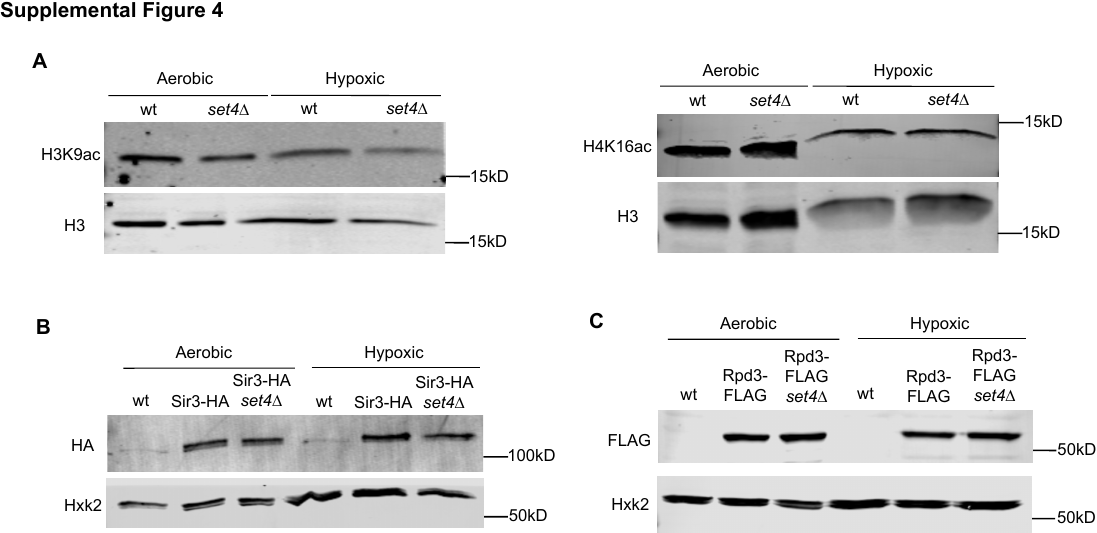

**Figure S4: Global acetylation and HDAC levels in wt and *set4Δ* under aerobic and hypoxic conditions.**

**A.** Clarified whole cell extract for wt (yEG001) and *set4Δ* (yEG322) strains grown to mid-log phase in YPD under normal and hypoxic conditions. Immunoblots probed with antibodies against H3, H3K9ac and H4K16ac. **B**. Immunoblotting of Sir3-HA from clarified lysates of wt (yEG873) and *set4Δ* (yEG874) cells grown under aerobic and hypoxic conditions. Anti-Hxk2 immunoblot serves as a loading control. **C.** Immunoblotting of Rpd3-FLAG from clarified lysates of wt (yEG956) and *set4Δ* (yEG1010) cells grown under aerobic and hypoxic conditions.

Figure S5: Analysis of *sir2Δ* and *rpd3Δ* single mutants and double mutants in combination with *set4Δ*.

**A.** chIP of wt (yEG001) and Sir3-HA (yEG873) strains grown to mid-log phase in YPD in aerobic and hypoxic conditions. Percent input from at least three biological replicates is shown. **B, C.** Ten-fold serial dilutions of wt (yEG001), *set4Δ* (yEG322), *sir2Δ* (yEG917), *sir2Δset4Δ* (yEG997) and (**C**) wt (yEG919), *set4Δ* (yEG920), *rpd3Δ* (yEG921), *rpd3Δ set4Δ* (yEG922) spotted on YPD and grown in aerobic or hypoxic conditions. Images of aerobic plates were taken after 2 days of growth at 30°C and images of hypoxic plates were taken after 8 days of growth **D.** RT-qPCR of subtelomeric genes from wt (yEG001), *sir2Δ* (yEG917) and *rpd3Δ* (yEG921) strains grown under aerobic conditions in YPD. For all panels, the error bars indicate S.E.M. and asterisks represent *p* values as calculated by one way ANOVA and Tukey’s post-hoc test (* <0.05, **<0.01, *** <0.001).

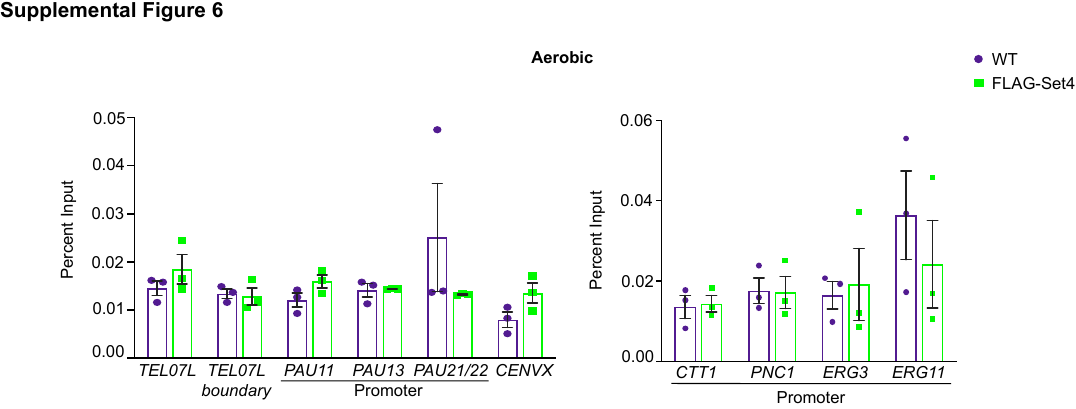

Figure S6: Set4 is not detectable at subtelomeric regions or stress response promoters under aerobic conditions.

chIP of FLAG-Set4 from cells grown under aerobic conditions. Percent input from three biological replicates is shown. Error bars represent S.E.M. and no significant differences were detected using an unpaired t test.
